# Allosteric Constraints on Rewiring Inducible Repressors

**DOI:** 10.64898/2026.08.06.743187

**Authors:** Abhilasha Gupta, Mitchell Lewis

## Abstract

Precise chemical control of transgene expression is central to synthetic biology, mammalian cell engineering, and gene therapy. Although tetracycline-responsive systems are widely used, converting an inducible repressor into a robust co-repressible regulator remains difficult. Systems engineered to activate DNA binding in response to ligand often exhibit elevated basal expression, weak switching, and limited dynamic range, suggesting that regulatory polarity is constrained by the underlying allosteric free-energy landscape.

Here we combine thermodynamic modeling with matched mammalian reporter assays to examine the fundamental distinction between inducible and co-repressible regulation. Using a promoter-occupancy framework, we describe how ligand binding redistributes regulators between DNA-binding–competent and DNA-binding–incompetent conformations to control transcriptional output. Inducible repressors such as TetR activate transcription by reducing operator occupancy, whereas co-repressible systems must increase operator occupancy to suppress transcription, imposing fundamentally different energetic requirements.

Experimental comparison of TetR-derived and PurR-derived regulators supports this thermodynamic interpretation. TetR-based systems produced strong ligand-dependent induction, whereas reverse TetR variants exhibited weaker co-repressible behavior and higher residual expression. In contrast, the natural co-repressible regulator PurR responded to hypoxanthine with ligand-stabilized DNA binding, and PurR–VP16 produced stronger ligand-dependent transcriptional activation than reverse TetR. Together, these results show that regulatory performance is determined by how efficiently ligand binding redistributes conformational states and suggest that natural co-repressible scaffolds may provide superior foundations for engineering ligand-activated transcriptional control.

## 1 Introduction

A central goal in gene regulatory engineering is to control transcriptional output in response to small molecules with high precision, reversibility, and tunability [1–3, 19]. Among the most widely used regulatory platforms are inducible repressors, in which ligand binding reduces DNA-binding affinity and relieves repression. These systems are powerful because they convert the presence of a small molecule into a quantitative and reversible change in gene expression.

Despite their broad utility, repression-based systems often suffer from elevated basal expression, particularly in mammalian cells. Because repressors frequently regulate relatively strong promoters, incomplete repression can produce substantial background transcription in the nominal OFF state. This residual leakiness limits applications requiring stringent control, including toxic transgene expression, developmental programs, and therapeutic gene delivery [18]. These limitations motivated the development of ligand-activated systems, in which transcription remains low in the absence of ligand and increases only upon induction. In such systems, weak or minimal promoters suppress basal transcription, while ligand binding promotes DNA occupancy and transcriptional activation, thereby separating low OFF-state expression from high induced output.

This raises a fundamental question: can an inducible repressor be rewired into a co-repressible or ligand-activated system while retaining comparable regulatory performance? Conceptually, such a transformation appears straightforward because inducible and co-repressible systems differ primarily in the direction of the ligand response. In inducible systems, ligand binding weakens DNA binding, whereas in co-repressible systems ligand binding strengthens DNA binding. In principle, converting one mode into the other should therefore require only inversion of the coupling between ligand binding and operator affinity. Indeed, engineered systems such as Tet-On demonstrate that ligand-dependent regulatory behavior can be experimentally reversed [4, 5].

In practice, however, reversing ligand-dependent regulation often compromises regulatory performance. Although engineered systems such as Tet-On successfully invert the direction of transcriptional control, they frequently exhibit substantially weaker switching behavior than the original inducible repressors from which they were derived. Common limitations include reduced dynamic range, elevated basal expression, incomplete repression or activation, and diminished ligand responsiveness [5, 9, 11, 18]. These observations indicate that polarity reversal requires more than simply modifying promoter context or appending activation domains.

At the molecular level, inducible and co-repressible regulation reflect opposite energetic preferences between DNA-binding–competent and DNA-binding–incompetent conformations. Ligand binding redistributes the equilibrium between these states, thereby altering promoter occupancy and gene expression. In TetR, doxycycline binding strongly stabilizes a DNA-binding–incompetent conformation, producing efficient dissociation from operator DNA and robust derepression [7, 8]. More generally, modern ensemble views of allostery emphasize that ligand binding changes the statistical population of pre-existing conformational states rather than acting through a single deterministic structural pathway [16]. Reversing regulatory polarity therefore requires inversion of the energetic coupling between ligand binding and DNA affinity, such that ligand binding instead stabilizes the DNA-binding state. However, engineered reverse TetR variants often undergo only partial redistribution between conformational states, compressing the achievable change in promoter occupancy and thereby reducing transcriptional dynamic range. Because these allosteric interactions have been evolutionarily optimized, inversion of regulatory polarity is energetically difficult and frequently weakens switching behavior.

Here, we examine the extent to which inducible repressors can be rewired into co-repressible and ligand-activated systems using a combination of thermodynamic modeling and experimental comparison. By analyzing TetR-derived regulators alongside naturally co-repressible transcription factors, we show that the principal limitation is not regulatory behavior itself, but the magnitude and directionality of allosteric coupling. These results suggest that, rather than forcing an inducible scaffold to adopt the opposite energetic response, it may be more effective to begin with regulatory proteins whose intrinsic allosteric properties already match the desired mode of control.

### 1.1 From Tet-Off to Tet-On: Reversing Regulatory Polarity

The first major adaptation of the bacterial tetracycline-resistance system for mammalian gene regulation was the Tet-Off system, in which TetR was fused to the potent herpes simplex viral activation domain VP16 to create the tetracycline-controlled transactivator (tTA) [6]. In the absence of doxycycline, tTA binds a tetracycline response element (TRE) containing tetO operator sequences positioned upstream of a minimal promoter and strongly activates transcription. Upon doxycycline binding, the equilibrium shifts toward a DNA-binding–incompetent conformation, causing dissociation from the operator and loss of transcriptional activation [7, 8]. Ligand addition therefore switches the system from an ON state to an OFF state.

Tet-Off demonstrated that a bacterial allosteric repressor could be successfully repurposed as a mammalian transcriptional activator capable of driving high levels of regulated gene expression [4, 6]. The system rapidly became a foundational platform for conditional gene expression because it combined strong transcriptional activation with tight operator specificity and reversible small-molecule control. However, because Tet-Off is intrinsically active in the absence of ligand, transcription must be continuously suppressed by maintaining doxycycline. For many biological and therapeutic applications, the opposite regulatory logic is preferable: minimal basal expression in the absence of drug and robust activation only after ligand addition [3, 9].

To reverse this ligand dependence, reverse TetR mutants were engineered that preferentially bind DNA in the presence rather than in the absence of doxycycline. These mutants formed the basis of the Tet-On system, in which doxycycline stabilizes the DNA-binding–competent state of the regulator and thereby promotes transcriptional activation [4, 5]. In contrast to Tet-Off, ligand binding therefore increases operator occupancy and activates transcription rather than disrupting DNA binding. Importantly, this transformation required more than appending an activation domain; it required rewiring the allosteric coupling between ligand binding and DNA-binding affinity.

Structural and biochemical studies of TetR have shown that doxycycline binding induces long-range conformational rearrangements that alter the relative positioning of the DNA-binding helices [7, 8, 10]. Converting TetR from an inducible repressor into a co-repressible or ligand-activated regulator therefore requires reversing the energetic linkage between ligand binding and DNA affinity. Such rewiring is intrinsically difficult because the conformational landscape of TetR evolved to destabilize DNA binding upon ligand association rather than promote it.

In practice, reversing allosteric response often compromises regulatory performance. Many Tet-On variants exhibit elevated basal expression, reduced ligand responsiveness, and weaker dynamic range than Tet-Off systems [5, 9, 11]. These observations suggest that the direction and efficiency of allosteric switching are not independently tunable, but are instead constrained by the underlying energetic properties of the regulatory protein. Together, these results imply that changing regulatory polarity is fundamentally an allosteric engineering problem rather than simply a promoter-design problem.

## 2 Experimental Strategy and Results

The preceding section framed polarity reversal as an allosteric engineering problem: ligand binding must not only change the direction of regulation, but must also redistribute the regulator strongly enough to produce a useful change in promoter occupancy. We therefore first tested this problem experimentally using matched Tet-derived systems, where Tet-Off provides a high-performing inducible reference point and Tet-On represents an engineered reversal of ligand response. We then used a thermodynamic occupancy model to interpret why reversal of polarity does not necessarily preserve dynamic range, before asking whether a naturally co-repressible regulator such as PurR provides a more favorable starting point for ligand-activated control.

### 2.1 Direct Comparison of Tet-Off and Tet-On Regulation

The Tet-Off and Tet-On systems provide a controlled experimental framework for examining whether regulatory polarity can be reversed without compromising switching performance [4, 6]. To compare these systems under matched conditions, we constructed autogenous regulatory modules in which a tetracycline response element (TRE) was positioned upstream of a minimal CMV promoter driving expression of luciferase together with either Tet-Off (TetR–VP16) or Tet-On (mutTetR–VP16 / rtTA). Since both circuits used the same promoter architecture, reporter, and activation domain, differences in output could be attributed primarily to ligand-dependent allosteric switching rather than to promoter topology or transcriptional effector composition.

Figure 1 shows luciferase expression measured in HEK293T cells in the absence and presence of doxycycline. Tet-Off displayed strong transcriptional activation in the absence of ligand and an approximately 16-fold decrease following doxycycline addition, consistent with efficient ligand-induced dissociation of TetR–VP16 from DNA [6]. In contrast, Tet-On exhibited only modest activation (∼1.3–1.8-fold) together with relatively elevated basal expression. Thus, although regulatory polarity was reversed, doxycycline produced only a limited increase in promoter occupancy and transcriptional activation.

**Figure 1:**
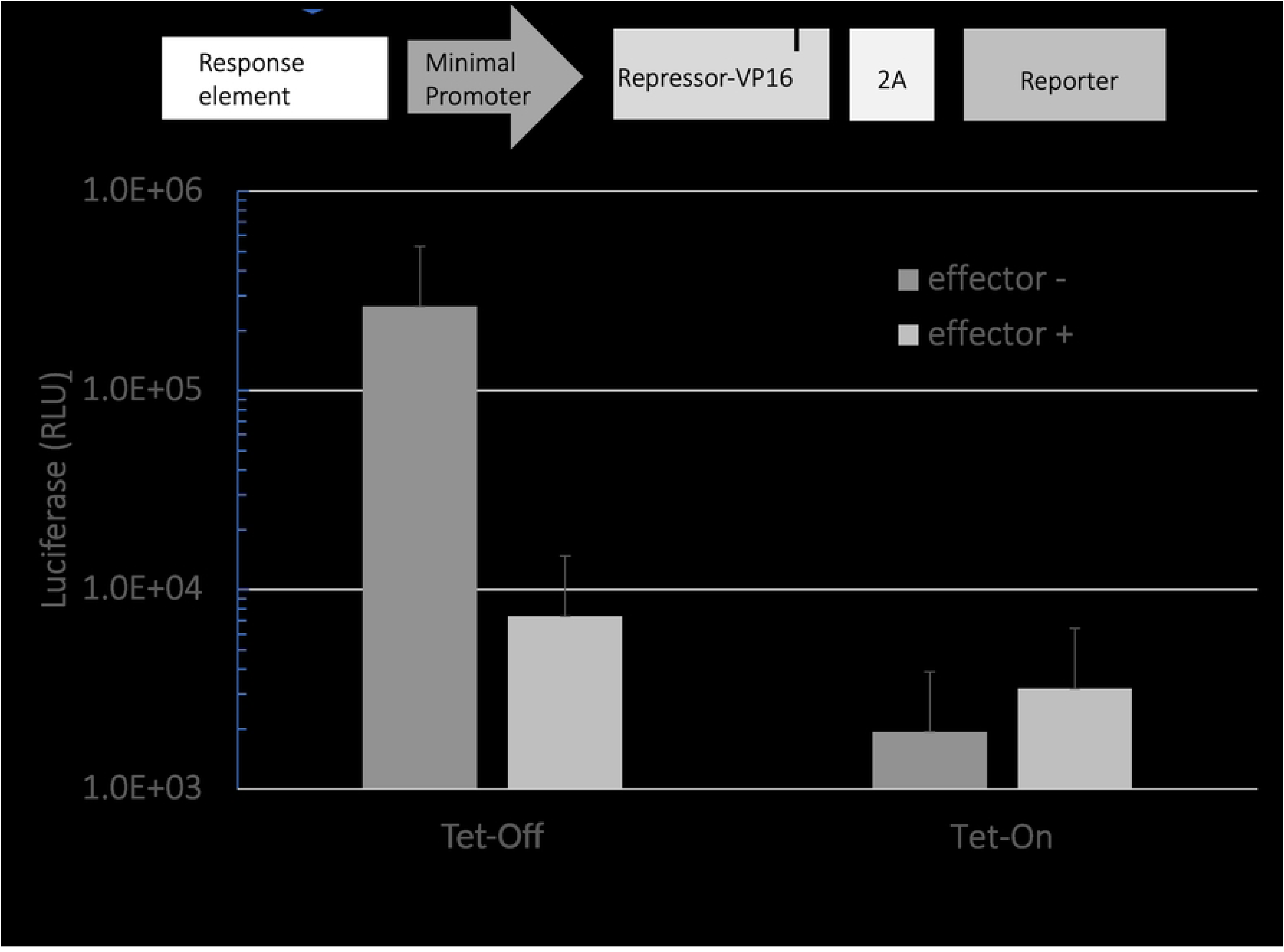
Comparison of Tet-Off and Tet-On transcriptional activators in a matched autogenous expression architecture. Top: schematic of the regulatory construct in which a tetracycline-responsive element (TRE) and minimal promoter drive expression of a repressor–VP16 fusion protein linked through a 2A peptide to a luciferase reporter. Bottom: luciferase expression measured in the absence (dark gray) or presence (light gray) of ligand. Tet-Off exhibited high expression in the absence of doxycycline and strong repression upon ligand addition, consistent with ligand-induced dissociation of TetR from DNA. Tet-On showed lower basal expression but only modest ligand-dependent activation. Data are shown on a logarithmic scale as mean ± SD from *n* = 6 replicate measurements.

These results show that inversion of regulatory logic is not equivalent to preservation of switching strength. Tet-On reverses the direction of ligand response, but the magnitude of ligand-dependent redistribution between DNA-binding–competent and DNA-binding–incompetent conformations is substantially weaker than in Tet-Off [5, 11]. This raised a second question: whether the weak Tet-On response reflects the VP16 activation domain and transcriptional activation context, or whether it reflects an intrinsic limitation of the engineered reverse TetR scaffold.

To separate these possibilities, we examined TetR and reverse TetR (rTetR) in a pure repression context lacking VP16. In these constructs, a CMV promoter containing tetO operator sites controlled expression of the repressor and a linked luciferase reporter. Because transcriptional output was determined solely by repressor occupancy, this design allowed ligand-dependent DNA binding to be evaluated independently of activation-domain function.

In the inducible CMV–TetR system, basal expression was low (∼ 1.0 × 10^5^ RLU) and increased to ∼ 1.4×10^6^ RLU following doxycycline addition, corresponding to approximately 14-fold induction (Fig. 2). By contrast, the co-repressible CMV–rTetR system exhibited higher basal expression (∼ 2.0 × 10^5^ RLU) that decreased only to ∼ 6.0 × 10^4^ RLU after ligand addition, corresponding to approximately 3-fold repression. Thus, even in the absence of VP16, rTetR displayed substantially weaker switching behavior than TetR.

**Figure 2:**
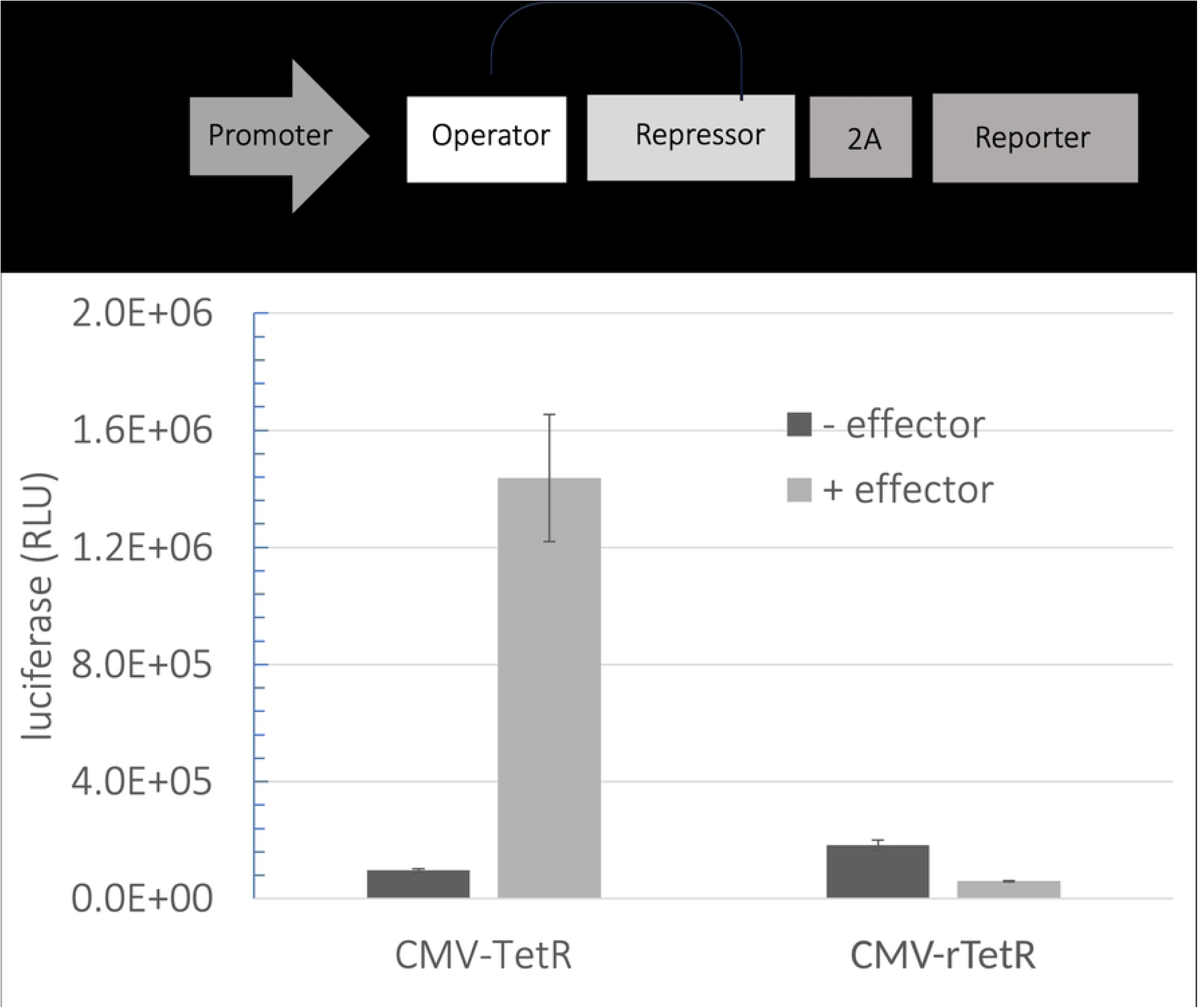
Autogenous repression circuits comparing inducible and co-repressible TetR regulators Top: schematic of the autogenous regulatory module in which a promoter containing an operator sequence drives expression of the repressor and a linked luciferase reporter through a 2A peptide. Bottom: luciferase output for the inducible TetR circuit (CMV–TetR) and the engineered co-repressible reverse TetR circuit (CMV–rTetR), measured in the absence (dark gray) or presence (light gray) of effector ligand. Data are shown as mean ± SD from *n* = 6 replicate measurements.

Together, the activation and repression experiments point to the same conclusion: the limited dynamic range of Tet-On systems does not arise primarily from the VP16 activation domain or from the transcriptional readout. Instead, it reflects weak ligand-dependent redistribution of DNA-binding occupancy by the engineered reverse TetR scaffold. This experimental result motivates a thermodynamic analysis of how conformational equilibria and ligand-binding preferences determine basal expression and dynamic range.

### 2.2 Thermodynamic Basis of Inducible and Co-repressible Regulation

The Tet experiments reveal a central mechanistic problem: reversing the direction of ligand response does not necessarily preserve the magnitude of regulatory switching. To understand this asymmetry, inducible and co-repressible systems can be described within a common statistical thermodynamic framework in which transcription is determined by promoter occupancy states [12–14, 17]. Because the experimental constructs above use autogenous architectures, promoter output and repressor abundance are self-coupled through negative feedback: the same promoter determines both reporter expression and production of the regulator that represses it.

In the simplest representation, the promoter exists in three mutually exclusive states: unbound, occupied by RNA polymerase (RNAP), or occupied by repressor. These states are assigned the statistical weights

| Promoter state | Weight |
| --- | --- |
| Empty | 1 |
| RNAP bound | $\alpha$ |
| Repressor bound | $\rho(L)$ |

where *α* represents the effective RNAP binding weight and

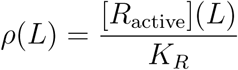

is the ligand-dependent repressor occupancy. Here, [*R*_active_](*L*) is the concentration of DNA-binding–competent repressor and *K_R_* is the dissociation constant for operator binding. The corresponding partition function is

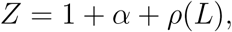

and transcriptional output is proportional to the probability of RNAP occupancy,

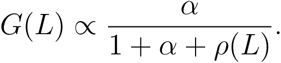

Ligand therefore regulates transcription indirectly by altering the population of repressor molecules that occupy the DNA-binding state.

The repressor is assumed to fluctuate between a DNA-binding–competent state (*R*) and a DNA-binding–incompetent state (*R*^∗^), with intrinsic conformational equilibrium

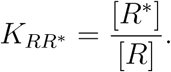

Ligand binds these conformations with different affinities, redistributing the conformational population and thereby altering DNA occupancy. Following the Monod–Wyman–Changeux (MWC) formalism [2, 15, 17], the fraction of repressor in the DNA-binding state for a dimeric regulator containing two ligand-binding sites is

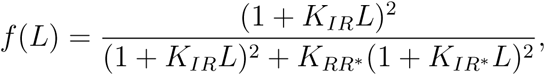

where *K_IR_*and *K_IR_*∗ are the ligand association constants for the DNA-binding and DNA-binding–incompetent conformations, respectively.

In an autogenous circuit, the repressor is produced from the same promoter that it regulates. Repressor abundance is therefore not fixed externally, but instead increases with promoter activity,

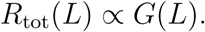

Because only a fraction *f* (*L*) of the repressor population is in the DNA-binding state, effective operator occupancy becomes

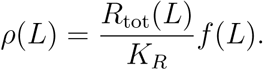

Defining a feedback parameter,

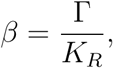

where Γ relates promoter output to repressor production, gives

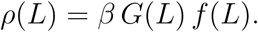

Substituting this expression into the promoter occupancy equation yields

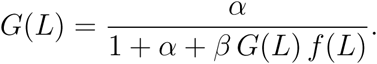

Thus, in an autogenous circuit, promoter activity and repressor abundance become mutually coupled: increased transcription produces more repressor, which in turn feeds back to suppress transcription. This feedback introduces a nonlinear constraint on gene expression. Within this framework, inducible and co-repressible systems differ in the direction of allosteric coupling. For inducible repressors such as TetR, ligand preferentially binds the DNA-binding–incompetent state,

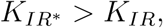

thereby decreasing operator occupancy and increasing transcription. Conversely, in co-repressible systems, ligand preferentially stabilizes the DNA-binding state,

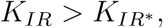

which increases operator occupancy and represses transcription.

The key thermodynamic parameter governing regulatory polarity is

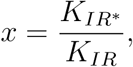

such that *x* ≫ 1 produces inducible behavior, *x* ≪ 1 produces co-repressible behavior, and *x* ≈ 1 yields weak ligand dependence. At saturating ligand concentrations, the effective conformational equilibrium scales approximately as

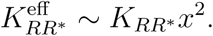

Thus, reversing regulatory polarity requires more than simply reversing the relative ligand affinities of the two conformational states,

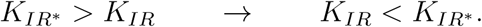

Robust co-repressible regulation additionally requires an intrinsic conformational equilibrium (*K_RR_*∗) that permits ligand binding to redistribute a substantial fraction of the regulator population into the DNA-binding state. If the underlying equilibrium strongly favors one conformation, reversing ligand preference alone produces only weak changes in promoter occupancy and therefore limited regulatory dynamic range.

In autogenous circuits, this molecular redistribution is further coupled to repressor abundance itself, because changes in promoter occupancy alter production of the regulator.

The induction profiles shown in Fig. 3 illustrate this thermodynamic constraint. The solid curve represents a classical inducible repressor such as TetR, in which ligand binding destabilizes the DNA-binding state. As ligand concentration increases, operator occupancy decreases sharply, producing efficient derepression.

**Figure 3:**
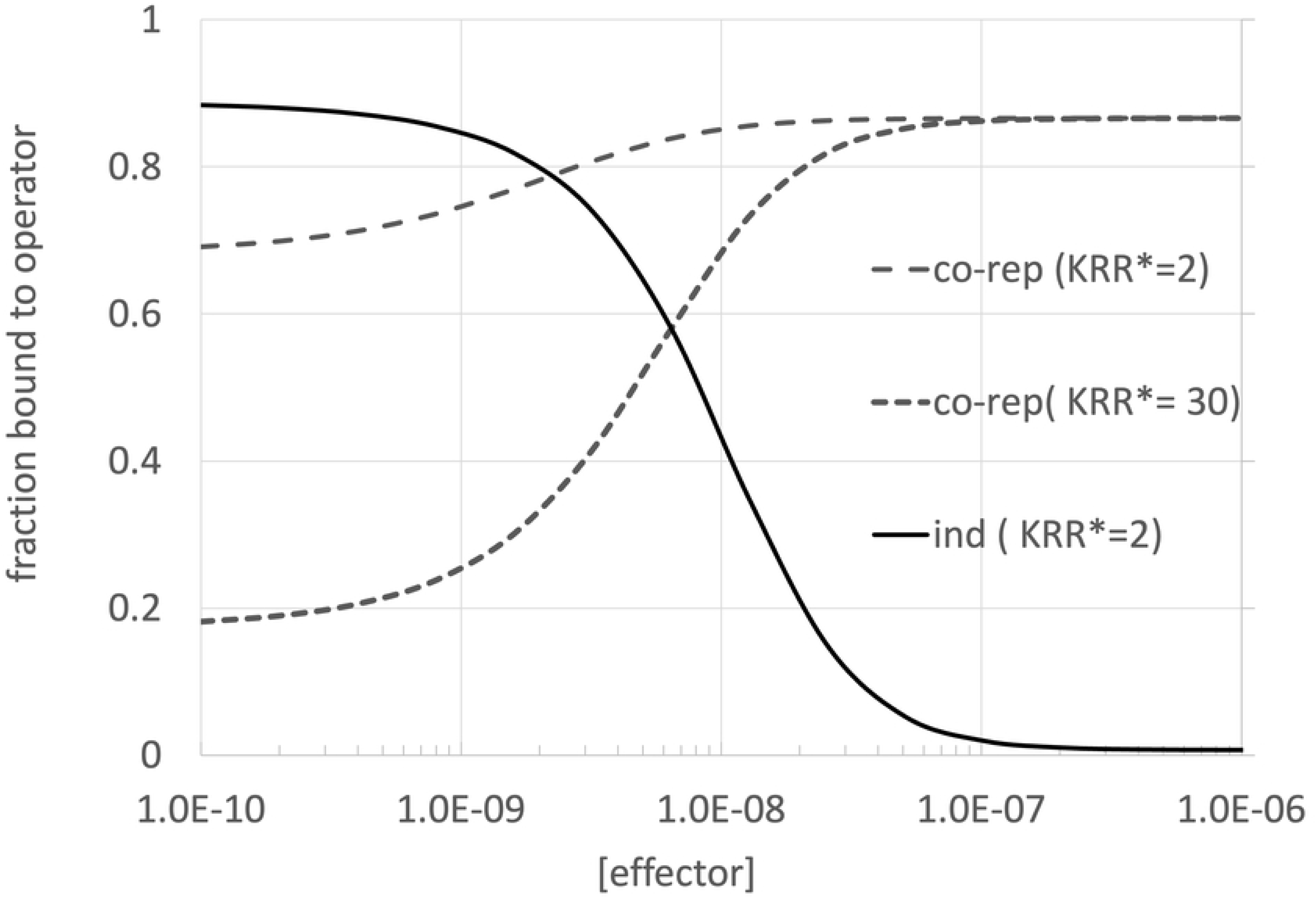
Thermodynamic model of inducible and co-repressible switching. The solid curve represents an inducible repressor, whereas the dashed and dotted curves represent co-repressible regulators with different intrinsic conformational equilibria (*K_RR_*∗).

The dashed and dotted curves instead represent co-repressible systems in which ligand binding stabilizes the DNA-binding state. Although both systems exhibit the same regulatory polarity, they differ substantially in basal occupancy and dynamic range because they possess different intrinsic conformational equilibria, described by the Monod linkage parameter *K_RR_*∗.

For the dashed curve (*K_RR_*∗ = 2), the equilibrium is only weakly biased toward the DNA-binding–incompetent state. As a result, a substantial fraction of regulator already occupies the DNA-binding state in the absence of ligand, producing high basal occupancy and only a modest ligand-dependent increase in DNA binding. By contrast, the dotted curve (*K_RR_*∗ = 30) strongly favors the DNA-binding–incompetent state without ligand, resulting in low basal occupancy but a much larger ligand-driven redistribution toward the DNA-binding state and therefore greater dynamic range.

These profiles illustrate a central thermodynamic principle originally emphasized by Monod and colleagues: regulatory behavior depends not only on differential ligand affinity between conformational states, but also on the intrinsic equilibrium between those states [2, 16, 17]. Reversing ligand-binding preference alone is therefore insufficient to produce robust co-repressible switching, because the underlying conformational equilibrium must also permit large occupancy redistribution.

The limited dynamic range of engineered reverse Tet systems therefore does not arise from VP16-mediated activation. Instead, both activation- and repression-based measurements support the same conclusion: inducible TetR undergoes strong ligand-dependent switching, whereas engineered co-repressible rTetR exhibits weaker allosteric coupling. The principal limitation of Tet-On systems thus originates from the underlying allosteric energy landscape rather than from the mode of transcriptional control. This observation motivates comparison with naturally co-repressible regulators, in which ligand binding intrinsically stabilizes the DNA-binding state.

### 2.3 PurR as a Co-repressible Scaffold

The limitations encountered in rewiring inducible repressors suggest an alternative strategy: begin with regulators that naturally exhibit co-repressible behavior. In these systems, ligand binding stabilizes the DNA-binding–competent conformation, increasing operator occupancy with increasing ligand concentration. This intrinsic allosteric coupling is already aligned with regulatory schemes in which ligand-dependent DNA binding modulates transcription.

PurR provides a prototypical example of a naturally co-repressible regulator [20]. In *E. coli*, PurR regulates genes involved in purine biosynthesis. Hypoxanthine and guanine act as co-repressors by stabilizing the DNA-binding state of PurR, thereby increasing binding to PurR operator sequences and repressing transcription. In the absence of co-repressor, PurR binds DNA more weakly and target genes are expressed. Thus, PurR is not simply an inverted inducible repressor; rather, its conformational equilibrium and ligand-binding energetics have evolved to support strong co-repression. As ligand concentration increases, the fraction of DNA-binding–competent regulator rises, producing a large and tunable increase in operator occupancy. Because this behavior arises from the intrinsic energetic properties of the protein, PurR does not require inversion of regulatory behavior to achieve ligand-dependent DNA binding.

### 2.4 Engineering a PurR-Based Transactivator

To determine whether a naturally co-repressible regulator could be repurposed for ligand-dependent transcriptional activation, we fused PurR to the VP16 activation domain. In this design, ligand binding increases the fraction of DNA-binding–competent PurR, which now recruits the transcriptional machinery and activates transcription. Thus, ligand-dependent changes in DNA occupancy are converted directly into ligand-dependent increases in gene expression.

Transcriptional output in the PurR–VP16 system depends not only on the fraction of DNA-binding–competent regulator, but also on the number and arrangement of operator sites within the promoter. To examine this dependence, we varied the number of PurR response elements positioned upstream of a minimal promoter and measured luciferase expression in HEK293T cells in the absence and presence of hypoxanthine.

Basal expression remained low across the operator series, near ∼ 10^3^ RLU, indicating minimal transcriptional activity in the absence of ligand when the DNA-binding state is weakly populated. Upon hypoxanthine addition, expression increased dramatically, and the magnitude of activation scaled strongly with operator copy number. With one response element, induced expression was approximately ∼ 10^4^ RLU. With two response elements, expression increased to ∼ 10^5^ RLU. With three response elements, expression approached ∼ 10^6^ RLU, and with four response elements, expression reached ∼ 10^7^ RLU. Because basal expression remained low while induced expression increased substantially, fold activation rose progressively with operator number, exceeding 10^3^-fold induction at the highest operator density (Fig. 4).

**Figure 4:**
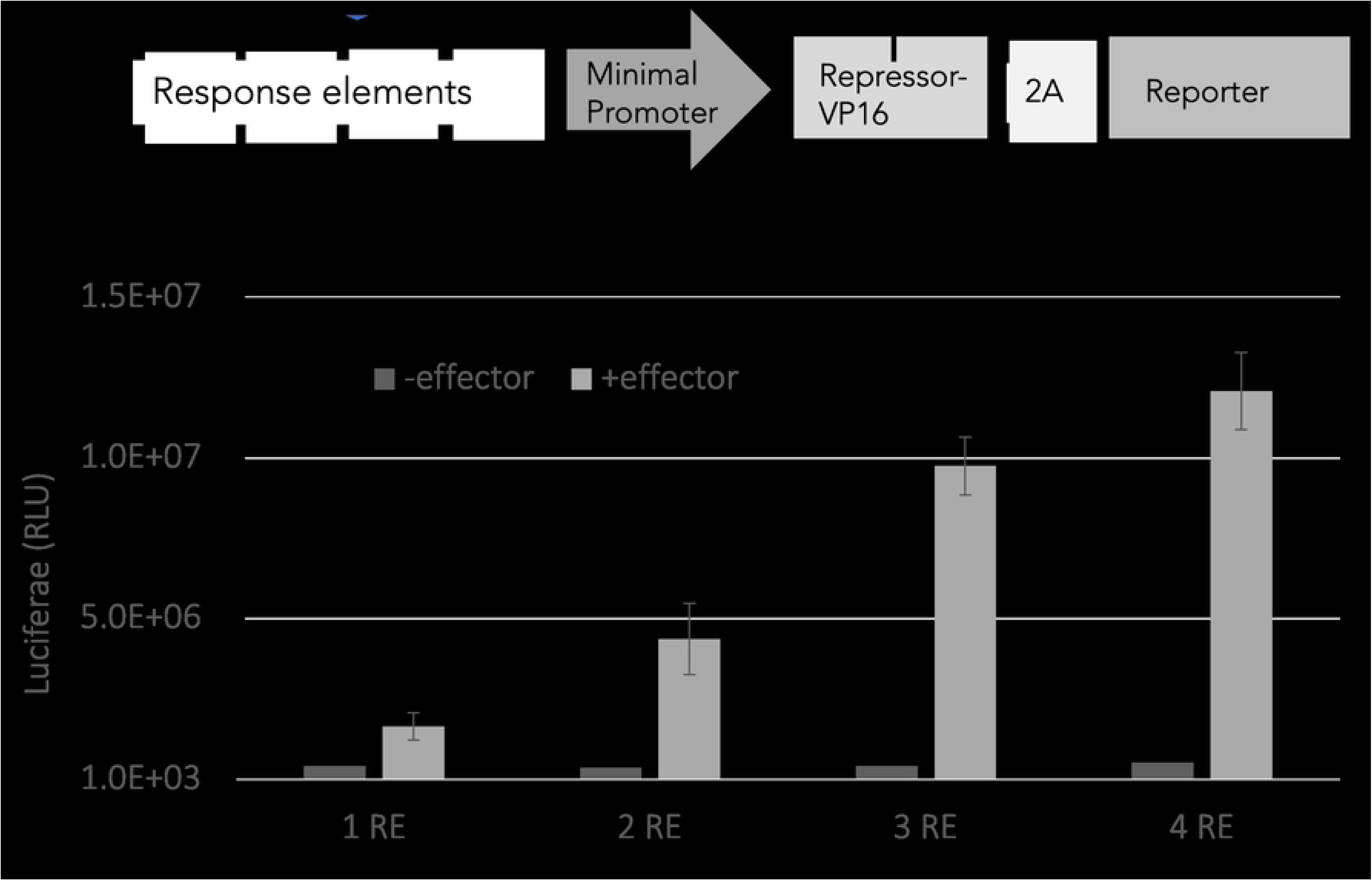
Effect of operator copy number on PurR–VP16 transcriptional activation. Top: schematic of the regulatory construct containing tandem response elements upstream of a minimal promoter driving expression of the PurR–VP16 activator and a linked luciferase reporter. Bottom: luciferase expression measured with one to four response elements (REs) in the absence (dark gray) or presence (light gray) of hypoxanthine. Data are shown as mean ± SD from *n* = 6 replicate measurements.

This behavior can be understood thermodynamically as an increase in the statistical weight of promoter states occupied by the activator. In a simplified description, the probability of transcription can be written as

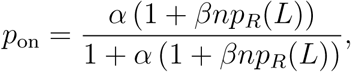

where *β* reflects the activation strength per bound regulator, *n* is the number of operator sites, and *p_R_*(*L*) is determined by the allosteric properties of PurR. In this framework, ligand controls the fraction of DNA-binding–competent regulator through *p_R_*(*L*), whereas promoter architecture scales transcriptional output through *n*.

Together, these results demonstrate that strong ligand-dependent allosteric coupling can be preserved while transcriptional output is tuned through promoter design. Natural co-repressors such as PurR therefore provide a favorable energetic foundation for constructing ligand-activated gene expression systems with both low basal expression and large dynamic range.

### 2.5 Comparison of Repressors and Transactivators

The regulatory systems shown in Fig. 5 reveal clear tradeoffs between basal expression, regulatory behavior, and dynamic range across inducible repressors, co-repressible repressors, and ligand-regulated transactivators. Together, these systems span nearly three orders of magnitude of expression and demonstrate that circuit performance depends strongly on the underlying allosteric mechanism.

**Figure 5:**
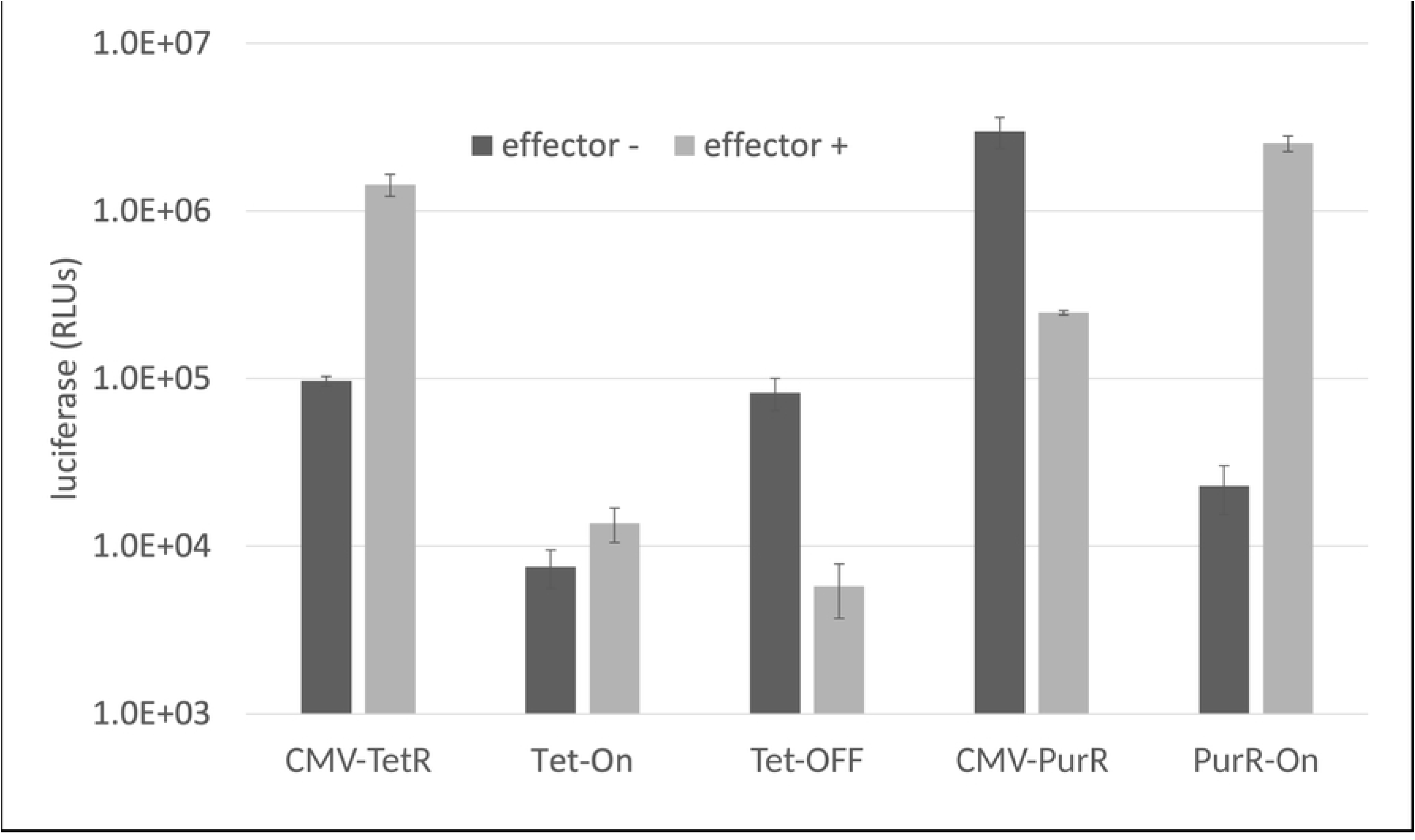
Comparison of TetR- and PurR-based transactivators in HEK293T cells. Tet-On (rtTA) exhibited low basal expression and only modest doxycycline-dependent activation, whereas Tet-Off (tTA) showed strong expression in the absence of ligand and repression following doxycycline addition. In contrast, PurR–VP16 combined low basal expression with strong hypoxanthine-dependent activation, producing the largest dynamic range among the systems tested. These results demonstrate that preserving a naturally co-repressible allosteric response enables stronger ligand-dependent activation than engineered reversal of TetR polarity. Bars represent mean ± SD from *n* = 6 replicate measurements.

Among the repression-based systems, the inducible TetR repressor (CMV-TetR) demonstrated robust switching behavior with relatively low basal expression. In the absence of doxycycline, expression remained near ∼ 1 × 10^5^ RLU, indicating efficient repression by operator-bound TetR. Addition of doxycycline relieved repression and increased expression to approximately 1.5 × 10^6^ RLU, corresponding to roughly 15-fold induction. TetR therefore combines low OFF-state leakiness with high induced output, producing the large dynamic range expected of an efficient inducible repressor.

The Tet-Off transactivator also exhibited strong switching, but with inverted regulatory behavior. In this system, transcription was high in the absence of ligand (∼ 8 × 10^4^ RLU) because TetR–VP16 binds DNA and activates transcription without doxycycline. Ligand addition reduced expression to approximately 6 × 10^3^ RLU, corresponding to roughly 15-fold repression. Tet-Off therefore retains the strong allosteric-switching properties of TetR while coupling DNA binding to activation rather than repression. However, because expression is maximal in the absence of ligand, Tet-Off exhibits relatively high basal activity from the perspective of inducible gene activation.

In contrast, the engineered Tet-On transactivator achieved the desired regulatory behavior but exhibited substantially weaker switching. Basal expression in the absence of doxycycline was low (∼ 7 × 10^3^ RLU), but ligand addition increased expression only modestly to approximately 1.3 × 10^4^ RLU, corresponding to only ∼ 1.5–2-fold activation. Tet-On therefore produces ligand-dependent activation while maintaining low basal expression, but its dynamic range is severely compressed relative to TetR and Tet-Off.

The PurR repression system behaved differently because PurR functions as a naturally co-repressible regulator. In the absence of hypoxanthine, expression was very high (∼ 3 × 10^6^ RLU), reflecting weak DNA occupancy under ligand-free conditions. Addition of ligand strongly increased operator occupancy and reduced expression to approximately 2.5 × 10^5^ RLU, corresponding to roughly 10–15-fold repression. PurR therefore maintains very high maximal output while still producing strong ligand-dependent repression.

Most notably, the PurR–VP16 transactivator combined the favorable regulatory behavior of Tet-On with dramatically improved dynamic range. Basal expression remained relatively low (∼ 2 × 10^4^ RLU), yet hypoxanthine increased expression to nearly 3 × 10^6^ RLU, corresponding to greater than 100-fold activation and representing the largest fold induction among all constructs tested. Unlike Tet-On, PurR–VP16 retains the native co-repressible behavior of PurR while coupling DNA binding to transcriptional activation..

Taken together, these comparisons reveal a fundamental thermodynamic distinction between engineered polarity reversal and native co-repression. TetR naturally couples ligand binding to loss of DNA affinity, enabling strong inducible repression and robust Tet-Off switching. Reversing this allosteric response in Tet-On preserves low basal expression but substantially weakens ligand-dependent occupancy changes, thereby limiting activation. In contrast, PurR intrinsically couples ligand binding to stabilization of the DNA-binding state. As a result, PurR-based activators achieve both low basal expression and large ligand-dependent induction without requiring inversion of the underlying allosteric free-energy landscape.

### 2.6 Endogenous Effectors as a Source of Basal Expression

Despite its strong inducibility, PurR–VP16 exhibited higher basal luciferase expression relative to some TetR-derived regulators. We hypothesized that endogenous purine metabolites present in mammalian cells might act as co-repressors, partially stabilizing the DNA-binding conformation of PurR even in the absence of exogenous hypoxanthine. Such basal co-repressor occupancy would increase promoter occupancy and elevate background transcriptional activation in the PurR–VP16 system.

To test this hypothesis, we examined substitutions at residue T192, a key determinant of hypoxanthine-mediated activation [20]. The wild-type PurR–VP16 construct exhibited relatively high basal expression of approximately 7 × 10^4^ RLU. In contrast, all three T192 variants showed substantially reduced basal activity (Fig. 6). The T192D mutant reduced basal expression to approximately 1.5–2.0 × 10^4^ RLU, whereas T192E and T192Q further lowered basal activity to approximately 8 × 10^3^–1.0 × 10^4^ RLU. Thus, substitutions at this position strongly suppressed background activation, consistent with reduced co-repressor binding and a shift of the conformational equilibrium toward the DNA-unbound state.

**Figure 6:**
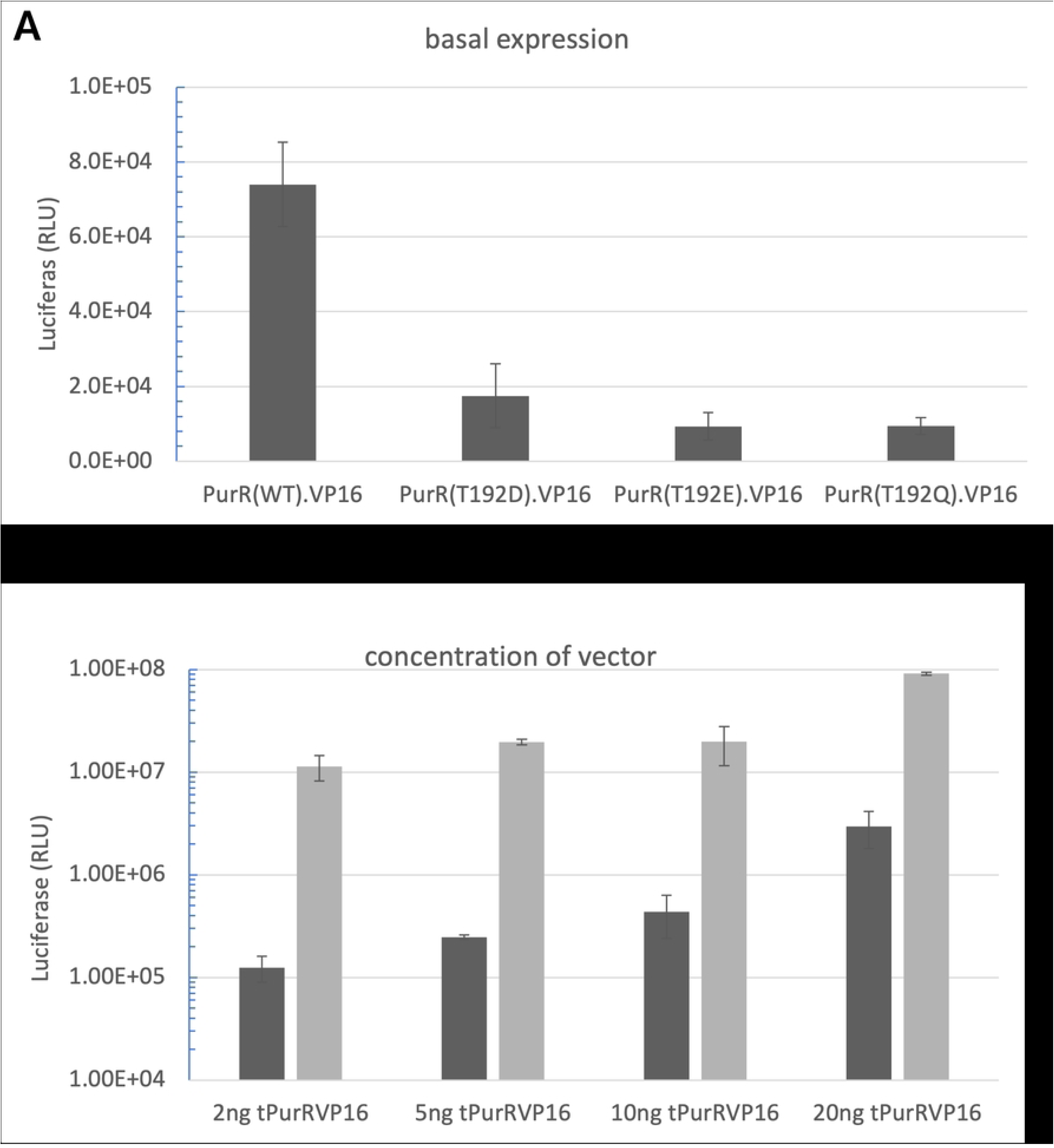
Basal expression and vector concentration analysis of PurR–VP16 variants. Top: basal expression of PurR(WT)–VP16 and T192 variants (T192D, T192E, T192Q), showing reduced activity in all mutants relative to wild type. Bottom: vector dose-dependent basal expression of PurR–VP16 at 2, 5, 10, and 20 ng, demonstrating progressive increases in signal with increasing vector input. Bars represent mean ± SD from *n* = 6 replicate measurements.

The reduction in basal activity produced by mutation of T192 supports the model that endogenous purine metabolites contribute to background activation in the wild-type construct. Disrupting co-repressor binding reduces the fraction of DNA-binding–competent regulator in the absence of added ligand, thereby decreasing promoter occupancy and lowering transcriptional output.

Basal expression was also strongly dependent on vector dosage. At 2 ng of vector input, basal activity was approximately 1.2 × 10^5^ RLU. Increasing the dose to 5 ng raised basal expression to ∼ 2.5×10^5^ RLU, whereas 10 ng produced a further increase to ∼ 4–5×10^5^ RLU. At 20 ng, basal expression increased sharply to approximately ∼ 3×10^6^ RLU, representing an approximately 25-fold increase relative to 2 ng. The gradual increase observed at lower vector doses, together with the sharp rise at high vector concentrations, indicates that regulator over expression substantially elevates background activity.

Together, these results demonstrate that basal expression in the PurR–VP16 system is determined by both endogenous ligand occupancy and regulator abundance. Mutations that weaken co-repressor binding reduce basal activity, whereas increasing vector dosage amplifies it. Optimizing both parameters is therefore critical for minimizing background expression and maximizing ligand-dependent dynamic range. We hypothesize that basal expression could be reduced by genetically modifying the PurR co-repressor binding pocket to preferentially recognize allopurinol, a structural analog of hypoxanthine.

## 3 Discussion

The results presented here demonstrate that inducible and co-repressible gene regulation are governed by a common thermodynamic framework, yet differ fundamentally in how ligand binding redistributes regulator conformational states. Across TetR-, rtTA-, and PurR-based systems, dynamic range was determined not simply by ligand-binding affinity or regulatory behavior, but by the extent of ligand-dependent redistribution between DNA-binding–competent and DNA-binding–incompetent states. Together, these findings reveal a thermo-dynamic asymmetry between inducible and co-repressible regulation.

A central conclusion of this study is that regulatory behavior is thermodynamically asymmetric. In inducible repressors such as TetR, ligand binding stabilizes the DNA-binding–incompetent state, reducing operator occupancy and relieving repression. In co-repressible systems, by contrast, ligand binding stabilizes the DNA-binding–competent state, increasing promoter occupancy and repression. Although these mechanisms appear superficially symmetric, our results demonstrate that reversing ligand-dependent regulation while preserving strong switching behavior is intrinsically difficult.

This asymmetry is illustrated most clearly by the comparison between Tet-Off and Tet-On. Tet-On was engineered so that doxycycline stabilizes the DNA-binding state rather than the DNA-binding–incompetent state. Although this modification successfully reversed regulatory behavior, the resulting system exhibited only weak ligand-dependent activation compared with the strong switching observed in Tet-Off. Similar behavior was observed in repression-based comparisons between TetR and rTetR, demonstrating that the limited dynamic range of Tet-On does not arise from VP16-mediated activation alone. Instead, reduced performance reflects incomplete redistribution of conformational occupancy within the engineered allosteric landscape.

These observations emphasize that dynamic range depends not simply on ligand affinity, but on how effectively ligand binding redistributes the regulator population between DNA-binding–competent and DNA-binding–incompetent states. Effective co-repressible regulation therefore requires both differential ligand binding and an intrinsic conformational equilibrium capable of supporting large ligand-dependent occupancy shifts. If ligand binding alters conformational occupancy only modestly, promoter occupancy and transcriptional output will likewise change only modestly, even when regulatory behavior has been successfully reversed.

The experimental data further demonstrate that repressor architecture amplifies these molecular limitations. Repression- and activation-based systems operate in fundamentally different promoter regimes and therefore experience distinct constraints on basal and maximal expression. In repression-based systems, strong promoters favor RNA polymerase occupancy even in the presence of repressor. Consequently, incomplete repression produces elevated basal expression, and increasing promoter strength elevates both OFF- and ON-state output. Strong promoters therefore provide high maximal expression but intrinsically amplify leakiness.

Activation-based systems exhibit the opposite behavior. Minimal or weak promoters suppress basal transcription because RNA polymerase binding is intrinsically unfavorable in the absence of activation. However, weak promoters also limit maximal output unless ligand binding produces a sufficiently large increase in DNA occupancy and transcriptional recruitment. This limitation was particularly evident in Tet-On systems, where activation of weak promoters yielded only modest induction despite successful polarity reversal.

Importantly, weak promoter activation is not intrinsically limiting. When activation was driven by PurR–VP16, ligand-dependent induction became both strong and scalable. PurR-based activators maintained low basal expression while achieving large increases in transcriptional output, demonstrating that weak promoters can support robust activation when ligand binding produces large shifts in DNA-binding occupancy.

These findings identify natural co-repressors as favorable scaffolds for engineering ligand-activated gene regulation. Unlike Tet-On, which requires inversion of an inducible allosteric response, PurR already couples ligand binding to stabilization of the DNA-binding state. Engineering therefore modifies only the transcriptional interpretation of DNA binding while preserving the native regulatory logic of the protein. Fusion of PurR to VP16 converts ligand-dependent DNA binding directly into ligand-dependent transcriptional activation without requiring reorganization of the underlying allosteric landscape.

More broadly, these results suggest a general design principle for synthetic gene regulation [19]. Repressor architecture defines the operating constraints of a system, but achievable dynamic range is determined by the strength of allosteric coupling and the extent of ligand-dependent occupancy redistribution. Repression of strong promoters tends to amplify leakiness, whereas activation of weak promoters minimizes basal expression but requires strong allosteric switching to achieve high output. Effective engineering strategies should therefore begin with regulatory scaffolds whose intrinsic energetics already support the desired mode of control rather than attempting to reverse regulatory behavior through limited mutational changes.

Taken together, these findings demonstrate that inducible and co-repressible systems are not simply interchangeable regulatory architectures with opposite directionality. Instead, they reflect distinct thermodynamic landscapes that differ in how ligand binding redistributes conformational occupancy. Transforming an inducible repressor into a strong co-repressible regulator is therefore substantially more difficult than repurposing a natural co-repressor as a transcriptional activator. This thermodynamic asymmetry provides a mechanistic framework for understanding the limitations of current inducible systems and offers general principles for designing next-generation mammalian gene circuits.

## 4 Funding and Declarations

This work was funded by a Synergy Grant from the School of Medicine, University of Pennsylvania. The funder had no role in study design, data collection and analysis, decision to publish, or preparation of the manuscript.

### Data availability

All data generated or analyzed during this study are reported in the manuscript and its figure files. Raw data are avaliable DOI: 10.5061/dryad.ht76hdrzr and plasmid wshould be made available upon reasonable request from the corresponding author.

### Author Contributions

Conceptualization: Mitchell Lewis.

Methodology: Mitchell Lewis, Abhilasha Gupta.

Investigation: Abhilasha Gupta.

Formal analysis: Mitchell Lewis, Abhilasha Gupta.

Writing – original draft: Mitchell Lewis.

Writing – review & editing: Mitchell Lewis, Abhilasha

Gupta. Funding acquisition: Mitchell Lewis.

Supervision: Mitchell Lewis.

## 5 Materials and Methods

### 5.1 Plasmid Design and General Cloning Strategy

All plasmid constructs used in this study were derived from the previously described autogenously regulated expression system (ARES) developed for gene-therapeutic applications [21]. The parental plasmid, pSW2.Luc, contained (from the 5^′^ to 3^′^ direction) an AAV2 ITR, a CMV immediate-early enhancer/promoter containing a symmetrical lac operator between the TATA box and transcription start site, a synthetic intron (Promega), the repressor coding sequence, a P2A cleavage site, firefly luciferase (Luc2; Promega), an SV40 polyadenylation signal, and the opposing AAV2 ITR. The promoter and operator region therefore controlled expression of both the regulator and luciferase, preserving the autogenous feedback architecture analyzed in the thermodynamic model. The principal LacR construct, pSW.CMV.2xOsym.LacR.ffLuc, contained two symmetrical operators rather than the single operator in pSW2.Luc.

Individual functional elements were flanked by unique restriction endonuclease sites, allowing promoters, operator arrays, DNA-binding proteins, activation domains, and secondary regulatory elements to be exchanged without reconstructing the complete vector. The regulatory cassette was positioned between AAV2 inverted terminal repeats (ITRs) to maintain compatibility with recombinant adeno-associated viral (AAV) vector production. Synthetic double-stranded DNA fragments (gBlocks; Integrated DNA Technologies) were used for most sequence replacements and operator-array assemblies. Inserts and vector backbones were digested with the indicated restriction enzymes, ligated in the required orientation, and propagated in *Escherichia coli*. The complete sequence of each modified region, including cloning junctions, was verified on both strands before the plasmid was used experimentally.

For TetR ARES constructs, the lacI regulatory region was excised with SacI and SpeI and replaced with a gBlock containing a CMV promoter with two tet operators separated by a 2-bp spacer, the TetR coding sequence, and a synthetic intron. Additional LacR and TetR variants were generated by replacing operator arrays, operator spacing, promoter length, or secondary regulatory elements with gBlocks flanked by compatible restriction sites.

### 5.2 Construction of Tet-On and Tet-Off Transactivators

The Tet-On transactivator construct was generated by PCR amplification of a 593-bp fragment containing the tetracycline response element (TRE), minimal CMV promoter, and multiple-cloning site from pTRE2 (Takara Bio/Clontech, cat. no. 631008). This fragment was joined to the rtTA–VP16 coding sequence to produce a 1,492-bp NheI–SpeI fragment, which replaced the corresponding region of pSW2.Luc to generate pSW.rtTA.Luc. The construct was subsequently modified so that the StuI site and Kozak sequence matched the repressor constructs. The TRE provides rtTA-binding sites, whereas the minimal CMV promoter limits transcription in the absence of bound transactivator.

The matched Tet-Off construct was produced from the Tet-On backbone by replacing reverse TetR with the conventional TetR–VP16 tetracycline-controlled transactivator using StuI and SpeI. Retaining the same promoter, reporter, activation domain, and vector context allowed the two constructs to differ principally in the polarity of their TetR-derived allosteric response. The VP16 activation domain, derived from herpes simplex virus VP16, recruits the mammalian transcriptional machinery when the fusion protein occupies the TRE [23, 24].

### 5.3 Construction of PurR Corepressor and PurR–VP16 Circuits

The PurR coding sequence was derived from the *E. coli purR* gene originally described by Rolfes and Zalkin [25]. The coding sequence was optimized for mammalian expression and synthesized by ATUM in plasmid pJ201.PurR. PurR was then transferred into the autogenous expression backbone through the StuI and SpeI sites to generate a repression-based circuit lacking a heterologous activation domain.

PurR operator sequences were derived from previously characterized *purR* regulatory elements [26] and positioned upstream of the regulated promoter so that ligand-dependent PurR binding could directly alter promoter output. The complete AAV-compatible PurR corepressor genome, measured from the 5^′^ to the 3^′^ ITR, was approximately 4.3 kb.

For activation-based measurements, PurR was fused to the VP16 activation domain. VP16 was synthesized as a gBlock and inserted into the PurR plasmid through the MluI and SpeI sites, maintaining the PurR and VP16 sequences in frame. The resulting PurR–VP16 cassette replaced rtTA–VP16 in the minimal-promoter transactivator backbone. This design converted ligand-stabilized PurR occupancy into transcriptional activation while retaining the same luciferase readout used for the Tet-derived transactivators.

To test the effect of operator number, a series of otherwise matched constructs containing one to four tandem purO2 elements upstream of the minimal CMV promoter was assembled. Operator arrays were synthesized as gBlocks with intervening linker sequences and inserted in the same promoter position and orientation. Consequently, changes in reporter output across the series could be attributed to operator copy number within a shared vector and promoter context.

For the principal PurR repression construct, the first operator was placed 9 bp down-stream of the TATA box, followed by a 9-bp spacer and the second operator. The complete PurR cassette measured 4,274 bp between the 5^′^ and 3^′^ ITRs. The tPurR and tPurR–VP16 constructs used four sequential purO2 elements upstream of a minimal CMV promoter with linker sequences between operators.

Matched bi-directional constructs were generated from the ARES modules. In these constructs, a reverse-oriented promoter drove the repressor or transactivator, while the opposing operator–promoter cassette drove firefly luciferase; the two transcription units were separated by a transcriptional pause sequence. The PurR–VP16 bi-directional circuit used an hPGK promoter to drive PurR–VP16 in the reverse direction and a 4xpurO2–CMVmin cassette to drive luciferase in the forward direction. Related PurR, TetR, and LacR variants were generated by exchanging the repressor coding sequence and operator–promoter gBlocks. CMV enhancer/ minimal-promoter, CMV-IE, minP, TREGS, hPGK, and truncated VP16 configurations were evaluated to determine how promoter strength affected basal expression and switching.

### 5.4 Site-Directed Mutagenesis of PurR

PurR variants T192N, T192D, T192E, and T192Q, were generated using QuikChange Lightning site-directed mutagenesis (Agilent Technologies, cat. no. 210518). Complementary mutagenic primers were designed with the desired substitution near the center. For each 50-*µ*l reaction, 10 ng pJ201.PurR template was mixed with 10× reaction buffer, primer pairs (100 ng/*µ*l), dNTPs, QuikSolution, and Pfu Ultra polymerase. Cycling consisted of 95^◦^C for 1 min; 18 cycles of 95^◦^C for 50 s, 60^◦^C for 50 s, and 68^◦^C for 4 min; and a final 68^◦^C extension for 7 min. Products were checked for the expected 3,686-bp amplicon, digested with DpnI to remove parental template, and transformed into XL10-Gold ultracompetent cells using NZY^+^ recovery medium (SOC was used as an acceptable substitute).

Candidate clones were screened by sequencing across the complete mutagenized region. Sequence-confirmed variants were subsequently transferred into both the PurR corepressor and PurR–VP16 expression vectors through the MluI and SpeI sites. This approach allowed each substitution to be compared in repression- and activation-based architectures without altering the surrounding promoter or reporter elements.

### 5.5 Cell Culture

HEK293T,cells were obtained from ATCC (Manassas, VA, USA), Cells were cultured in high-glucose DMEM H-glu GLUTAMAX with 10% fetal bovine serum, whereas ARPE-19 cells were cultured in DMEM/F-12 with 10% fetal bovine serum. All media contained 1% penicillin/streptomycin. Cultures were maintained at 37^◦^C in a humidified incubator containing 5% CO_2_ and were expanded under standard adherent-cell conditions before plating for transfection or transduction experiments. Cells used for reporter assays were plated at the densities specified below so that the different cell types reached an appropriate density at the time of nucleic-acid delivery.

### 5.6 AAV Vector Production and Transduction

Recombinant AAV8 vectors were produced by the CAROT Research Vector Core at the University of Pennsylvania as previously described (Duong et al., 2019). Vector genomes containing the regulatory cassette between AAV2 ITRs were packaged in AAV8 capsids following production in HEK293 cells. Viral particles were purified by CsCl ultracentrifugation. The AAV panel included LacR, TetR, TetR–DHFR, rtTA, and PurR–VP16 vectors, including both repressor and transactivator configurations and the corresponding bi-directional variants.

For transduction experiments, 84-31, HEK293T-AAVR, and ARPE-19-AAVR cells were seeded at 4 × 10^4^ cells per well in 100 *µ*l medium and exposed at seeding to 10^4^, 10^5^, or 10^6^ vector genomes per cell. Luciferase activity was measured at 24, 48, and 72 h after vector addition to follow the development of transgene expression over time.

### 5.7 Induction Conditions

Small-molecule treatments were initiated 4 h after transfection or transduction. I tetracycline, doxycycline, hypoxanthine, or allopurinol was added as appropriate for the regulatory system under study. Matched untreated wells were included to define basal reporter activity. Aptazyme two-component systems were treated with the relevant inducer plus tetracycline; DHFR dual switches received the relevant inducer plus trimethoprim to stabilize DHFR; and three-component switches received the appropriate inducer, tetracycline, and trimethoprim. For dose-response experiments, ligand concentration was varied across wells and cells were incubated for a 20 h induction period at 37^◦^C before reporter measurement. Tet-derived constructs were evaluated with tetracycline or doxycycline, whereas PurR-derived constructs were evaluated with hypoxanthine or the indicated purine analog.

### 5.8 Luciferase Assays

Firefly luciferase activity was quantified with Bright-Glo Luciferase Assay Reagent (Promega). At the indicated endpoint, culture medium was removed and replaced with Bright-Glo reagent diluted 1:1 with phenol red-free RPMI medium. Luminescence was measured in the 96-well plate using an Infinite Tecan-200 plate reader operated with Tecan 2.0 software. The same reagent preparation and instrument settings were used for the matched conditions within each experiment.

Raw luminescence was reported as relative light units (RLU). Basal expression was defined as the signal measured in the absence of added ligand. For ligand-activated systems, fold induction was calculated as the induced signal divided by the basal signal. For ligand-repressed systems, the magnitude of repression was calculated by comparing the ligand-free signal with the signal measured after ligand addition. Where dose-response curves were analyzed, maximal and basal values refer to the upper and lower plateaus observed across the ligand series. Dynamic-range and fractional-expression measurements were normalized to the corresponding constitutive-promoter controls. Replicate numbers and the summary statistic used for each experiment are specified in the figure legends.

## 6 Acknowledgments

We thank the CAROT Research Vector Core at the University of Pennsylvania for AAV vector production.

